# miR-146a-3p packaged in small extracellular vesicles triggers fetal membrane inflammation in response to viral dsRNA through activation of Toll-like Receptor 7 and 8

**DOI:** 10.1101/2025.07.29.667498

**Authors:** Hanah M Georges, Abigail C Fischer, Paloma Casanova, Vikki M Abrahams

## Abstract

Maternal infection and chorioamnionitis are one of the leading causes of preterm birth and neonatal morbidity. The relationship and mechanisms linking bacterial infections and preterm labor are well researched, however, less is known about the mechanisms involved in how viral infections contribute to preterm labor. Previous work from our group demonstrated that following bacterial triggers, fetal membranes (FMs) express elevated miR-146a-3p which in turn acts as an intermediate danger signal by activating TLR8 to induce a robust inflammatory response. Using an established FM explant model system, the role of this and other TLR7/8-activating miRs in the propagation of viral-induced inflammation was investigated. Following exposure to the viral dsRNA mimic and TLR8 agonist, Poly(I:C), expression of FM tissue TLR7/8-activating miRs were not elevated. Despite this, FM secretion of pro-inflammatory IL-6 and IL-8 were increased in response to Poly(I:C) in a TLR7- and TLR8-dependent manner. To investigate alternative methods of miR delivery, small extracellular vesicles (sEVs) from FM supernatants were isolated and found to contain elevated levels of miR-146a-3p and miR-21a under Poly(I:C) conditions. Furthermore, Poly(I:C)-induced IL-6 and IL-8 responses were reduced in the presence of an inhibitor of sEV biogenesis/release, and IL-6 production was reduced in the presence of a miR-146a-3p inhibitor. Together, these data suggests that sEVs produced from virally-stimulated human FMs contain and deliver elevated miR-146a-3p which acts as a danger signal to drive perpetuate inflammation via TLR7 and TLR8 activation. This work demonstrates a novel and important role for sEV packaged TLR7/8 activating-miR-146a-3p in FM inflammatory responses to viral infections.

## Introduction

The pregnant population is particularly vulnerable to both bacterial and viral infections. However, maternal infection often goes undiagnosed and untreated, making it one of the largest contributors to maternal morbidity, preterm birth, and neonatal morbidity and mortality^1,2^. The contribution of bacterial infections to chorioamnionitis – inflammation of the fetal membranes (FMs) – and preterm birth, are well established^3^. However, most of our mechanistic knowledge about the effects of viral infections on pregnancy and preterm birth comes from studies on TORCH pathogens; infections which cross the placenta and infect the fetus^4^. However, TORCH pathogens represent a minor fraction of viral infections. With the more recent influenza H1N1 and SARS-CoV-2 pandemics, increased research has shown that viral infections and maternal immune activation can have negative impacts on pregnancy outcomes and on the developing fetus in the absence of vertical transmission^5,6^. A systemic maternal infection and maternal immune activation can contribute to placental inflammation, preterm birth, and developmental disorders^1,2,7^. Despite these connections, the mechanisms by which a maternal viral infection may lead to preterm birth are still largely unknown and remains a clinical concern.

The maternal-fetal interface has robust innate immune strategies in place to help protect against invading pathogens. For example, the fetal membranes (FMs) possess the ability to respond to pathogens through pathogen recognition receptors (PRRs)^8-10^, leading to an inflammatory response. Once activated, FM expressed PRRs, such as Toll like receptors (TLRs), can trigger inflammation through the production of pro-inflammatory cytokines and chemokines, which can then initiate the production of mediators of membrane weakening and the induction of labor^11,12^. In certain pregnancy pathologies, uncontrolled or untimely inflammation uses the same pathways as normal term labor, resulting in preterm birth^13-15^.

Using a model of bacterial infection, our group recently reported a role for endogenous danger signals or DAMPs (damage associated molecular patterns) as intermediate mediators of FM inflammation downstream of an initial TLR sensor^12,16,17^. We found that FMs treated with bacterial triggers expressed elevated levels of a TLR7/8-activating microRNA^18^, miR-146a-3p^19,20^, which downstream of TLR2 and TLR4 activation by bacteria peptidoglycan (PDG) and lipopolysaccharide (LPS), respectively, drives FM inflammation and the production of mediators of membrane weakening through activation of TLR8^12,16^. The identification of such intermediate mediators of inflammation provides a new fundamental understanding of the underlying mechanisms of chorioamnionitis and preterm birth.

While previous bacterial models focused on locally tissue produced TLR7/8-activating miRs as mediators of inflammation, there is potential for alternate delivery methods of pro-inflammatory DAMPs. One potential source of intermediate inflammatory signals are small extracellular vesicles (sEVs) which provide cells the ability to communicate in both autocrine and paracrine manners, and are now known to be critical conduits of cellular communication^21^. Indeed, TLR7/8-activating miRs can be released via sEVs and delivered to target cells to exert a stimulatory effect^18,19,22^.

Using a well-established human FM explant model^12,16^, we investigated the roles of tissue expressed and sEV-associated TLR7/8-activating miRs as mechanisms for driving FM inflammation in response to a double stranded RNA (dsRNA) viral mimic, Poly(I:C). This work reveals an important role for FM-derived sEVs in the delivery of TLR7/8-activating miRs and subsequent inflammatory signaling in response to a viral trigger.

## Materials and Methods

### Patient samples

Human tissue collections were approved by Yale University’s Human Research Protection Program (IRB# 0607001625). Patients undergoing uncomplicated term (38-41 weeks) elective cesarean sections without labor or known infection were consented and FMs were collected by the Yale University Reproductive Sciences (YURS) Biobank (IRB# 1309012696) immediately after delivery. FMs were biopsied and cultured as previously described^12,16^. Briefly, FMs were rinsed, biopsied with a 6mm punch, and placed into 0.4μm cell culture inserts in a 24 well plate (BD Falcon, Franklin Lakes, NJ) containing 1mL of Dulbecco modified eagle medium (DMEM; Gibco, Grand Island, NY) supplemented with 10% fetal bovine serum (Hyclone, Logan, UT) and 1% penicillin streptomycin (Gibco). Explants were incubated overnight, then media was replaced with serum-free OptiMEM (Gibco), rested for 3 hours, and treated.

### Fetal membrane explant treatments

Human FM explants were treated with either no treatment (NT) as a media control or with 20μg/ml of high molecular weight Poly(I:C) (Invivogen; San Diego, CA) for 24 hours. We previously reported that at this dose Poly (I:C) triggers an inflammatory cytokine/chemokine response^23^. For inhibitor studies, human FMs were pretreated for 1 hour with either with media, the TLR7 inhibitor, IRS661 (5μM; 5′-TGCTTGCAAGCTTGCAAGCA-3’; made by the Keck Core, Yale University)^16^, the TLR8 inhibitor, CU-CPT9a (10μM; Invivogen); or the inhibitor of sEV biogenesis/release, GW4869 (0.1μM; R&D Systems; Minneapolis, MN). In other experiments, FM explants were transfected using siPORT NeoFX (Invitrogen, Waltham, MA) with either a scramble control, the miRVana inhibitor to miR-146a3p, or the miRVana inhibitor to miR-21a, all at 100nM (ThermoFisher Scientific, Waltham, MA) for 24 hours as previously described^16^, prior to NT or Poly(I:C) treatments. After 24 hours, FM supernatants and explants were collected and stored at -80°C for subsequent analyses.

### FM cytokine/chemokine measurements

Human FM explant supernatants were collected and centrifuged to clear the supernatants of debris. Supernatants were measured for known markers of inflammation/chorioamnionitis^16^: IL-6, IL-1β, and IL-8 by ELISA (R&D Systems, Minneapolis, MN) according to manufacturer’s protocol. Absorbance was measured using an iMark microplate absorbance reader (Bio-Rad) and reported as pg/ml.

### Small extracellular vesicle isolation and validation

Small extracelllular vesicles (sEVs) were isolated from FM supernatants using ExoQuick-TC (Systems Biosciences LLC, Palo Alto, CA) per the manufacturer’s instructions. The isolated sEV pellet was either: 1) resuspended in 50μl of phosphate buffered saline (PBS, Thermo Fisher Scientific, Waltham, MA) for subsequent nanoparticle tracking analysis and quantification using a Zetaview (Particle Metrix, Ammersee, Germany) set to room temperature, a laser wavelength of 488nm, a sensitivity of 80, and a shutter speed of 100; 2) directly lysed for RNA extraction using the using the SeraMir Exosome RNA Purification Column Kit (Systems Biosciences) as previously described^22^; or 3) directly lysed into sample buffer for Western blot analysis of the surface protein marker CD9 (#13174, Cell Signaling Technology; Danvers, MA) and CD81 (#56039T, Cell Signaling Technology).

### RT-qPCR

RNA was isolated from FM tissue explants as previously described^17^. Tissue and sEV RNA concentrations were measured with a NanoDrop 2000 Microvolume Spectrophotometer (ThermoFisher Scientific; Waltham, MA). Reverse transcription and qPCR to measure the expression of miR-21a, miR-29a, miR-146a-3p, and Let-7b was performed using the Taqman MicroRNA Assay (Life Technologies; Carlsbad, CA) according to manufacturer’s protocols. For miR-146a-3p, a pre-amplification step was included prior to performing PCR using the Taqman PreAmp master mix kit (Life Technologies). Small nuclear RNA U6 was a reference gene for normalization and data were analyzed with the 2^-ΔΔCT^ method and reported as fold change (FC).

### Statistics

For each human FM sample, explant treatments were performed in 3 technical replicates, pooled, and reported as one experiment. Experiments were performed in three or more biological/independent replicates and reported in the figure legends as “n=“. All analysis was performed in duplicate and averaged, and data are reported as mean ± SEM of independent experiments with statistical significance defined as *p*<0.05. Statistical analysis was performed using Prism Software (Graphpad, La Jolla, CA). Normality was determined using the Shapiro-Wilks test. For normally distributed data, data were analyzed using either one-way analysis of variance (ANOVA) for multiple comparisons or a t-test. For data not normally distributed, data were analyzed using a non-parametric multiple comparison test or the Wilcoxon matched-pairs signed rank test. When available, fetal sex was confirmed as an unsignificant variable as determined by multiple linear regression models. Additional statistical analyses are specified in table legends.

## Results

### Poly(I:C) increases FM tissue expression of TLR7/8-activating miRs

Previous work by our group has demonstrated that that elevated FM tissue expression of the TLR7/8-activating miR, miR-146a-3p, plays a role in mediating inflammation triggered by bacterial LPS or PDG^12,16^. To investigate the role of TLR7/8-activating miRs in a viral model, FM explants were exposed to a viral dsRNA mimic, Poly(I:C), and the tissues measured for the TLR7/8-activating miRs, miR-146a-3p, miR21a, miR-29a and Let-7b^18-20^. Following Poly(I:C) stimulation, expression levels of miR-146a-3p, miR-21a, and Let7b were all unchanged compared to the NT controls, and miR-29a levels were significantly decreased by 36.9±9.1% (Figure 1A).

**Figure 1.**
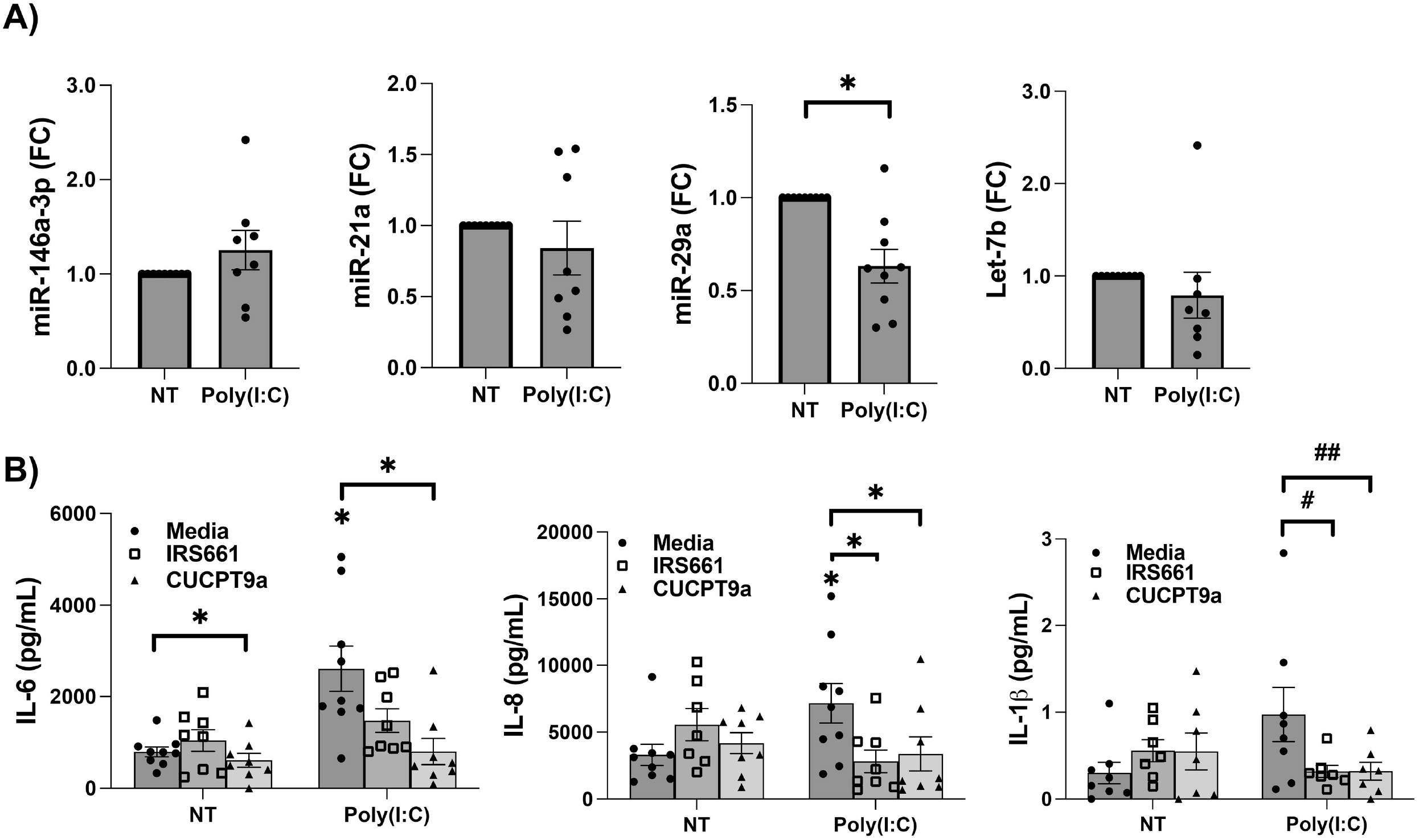
Poly(I:C) induces FM inflammation in a TLR7- and TLR8-dependent manner without elevating FM tissue expression of TLR7/8 activating microRNAs. (A) Human FM explants were either not treated (NT) as media controls or treated with Poly(I:C) for 24 hours and then tissues were collected for RNA extraction. The TLR7/8-activating miRs, miR-146a-3p, miR-21a, miR29a, and Let7b, were measured by RTq-PCR (n=8-9). ^*^*p*<0.05. (B) FM explants were pretreated with media, the TLR7 inhibitor, IRS661, or TLR8 inhibitor, CU-CPT9a, for 1 hour. FMs were then untreated (NT) or treated with Poly(I:C) and after 24 hour supernatants were measured for pro-inflammatory IL-6, IL-8, and IL-1β (n=8-9). ^*^*p*<0.05 relative to the NT/Media control unless otherwise indicated. #*p*=0.08; ##*p*=0.05.

### Poly(I:C) induces FM inflammation in a TLR7- and TLR8-dependent manner

Human FM explant stimulation with Poly(I:C) significantly increased the secretion of pro-inflammatory cytokine IL-6 and neutrophil recruiting chemokine IL-8 secretion by 3.4±0.6-fold and 2.2±0.3-fold, respectively (Figure 1B). While significance was not reached, FM secretion of pro-inflammatory cytokine IL-1β was increased by 6.8±2.1-fold in response to Poly(I:C) (Figure 1B). To investigate TLR7- and TLR8-dependent intermediate signaling previously reported in our bacterial models^12,16^, FMs were pre-treated with either a TLR7 inhibitor, IRS661, or a TLR8 inhibitor, CU-CPT9a. In Poly(I:C) stimulated FMs, inhibition of TLR7 significantly reduced FM IL-8 secretion by 45.0±18.6% compared to Poly(I:C) alone (Figure 1B). While not quite significant (*p*=0.08), inhibition of TLR7 reduced FM IL-1β secretion in response to Poly(I:C) by 12.9±42.1% (Figure 1B). TLR7 inhibition did not significantly alter Poly(I:C) induced FM IL-6 secretion (Figure 1B). In the presence of the TLR8 inhibitor, CU-CPT9a, Poly(I:C) stimulated FM secretion of IL-6 and IL-8 were both significantly reduced by 56.5±15.7% and 48.3±14.0%, respectively, when compared to Poly(I:C) alone (Figure 1B). TLR8 inhibition also decreased IL-1β secretion in Poly(I:C) treated FMs by 17.2±58.0%, however, this was not quite significant (*p*=0.05; Figure 1B).

### Poly(I:C) stimulated FMs release sEVs expressing elevated miR-21a and miR-146a-3p

With FM TLR3-mediated inflammation stimulated by Poly(I:C) being dependent on both TLR7 and TLR8, small extracellular vesicles (sEVs) were considered as an alternate method of TLR7/8-activating miR delivery. Thus, sEV release by FM explants and their miR cargo were investigated. The presence of sEVs in FM supernatants was confirmed by Western Blot for the markers CD81 and CD9 (Figure 2A). The presence of sEVs in FM supernatants was further validated using nanoparticle tracking analysis which confirmed sEV size (data not shown). As shown in Figure 2A, FMs released similar amounts of sEVs under both NT and Poly(I:C) conditions. Treatment of FMs with Poly(I:C) significantly elevated sEV expression of miR-146a-3p and miR-21a by 10.2±3.6-fold and by 2.8±0.7-fold, respectively (Figure 2B). sEVs from Poly(I:C) treated FMs expressed 4.8±1.6-fold more miR-29a when compared to the NT control, however, this was not quite significant (p=0.05, Figure 2B). Let7-b expression was unchanged in sEVs released from Poly(I:C) treated FMs when compared to no treatment (NT) controls (Figure 2B).

**Figure 2.**
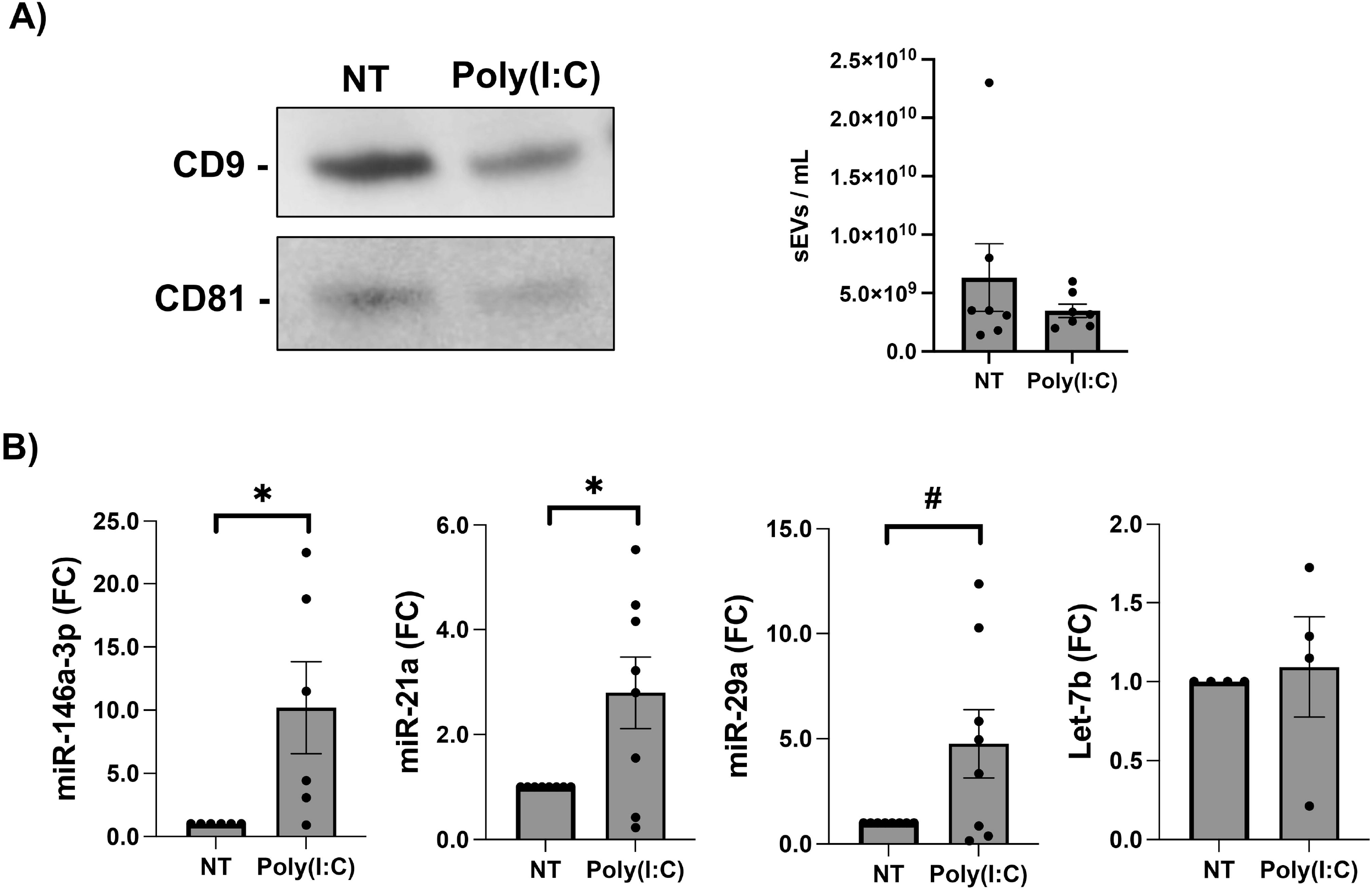
FMs treated with Poly(I:C) release sEVs containing elevated miR-146a-3p and miR-21a. (A) sEVs were isolated from the supernatants of FM explants treated for 24 hour with either NT or Poly(I:C). Image shows sEV expression of the surface marker CD9 and CD81 as determined by Western Blot. Chart shows sEV concentrations as determined by NTA. (B) RNA was isolated from sEVs and measured for the TLR7/8-activating miRs, miR-146a-3p, miR-21a, miR-29a and Let7b by RT-qPCR (n=4-8; ^*^*p*<0.05; #*p*=0.05).

### FM IL-6 and IL-8 secretion in response to Poly(I:C) is dependent on sEVs

The dependency of Poly(I:C) induced inflammation on sEVs was investigated using the inhibitor of sEV biogenesis/release, GW4869^24^. The presence of GW4869 significantly reduced the ability of Poly(I:C) to induce FM secretion of IL-6 and IL-8 by 23.6±9.4% and 25.1±19.1%, respectively (Figure 3A). In contrast, Poly(I:C)-induced FM IL-1β secretion was unchanged by sEV inhibition (Figure 3A).

**Figure 3.**
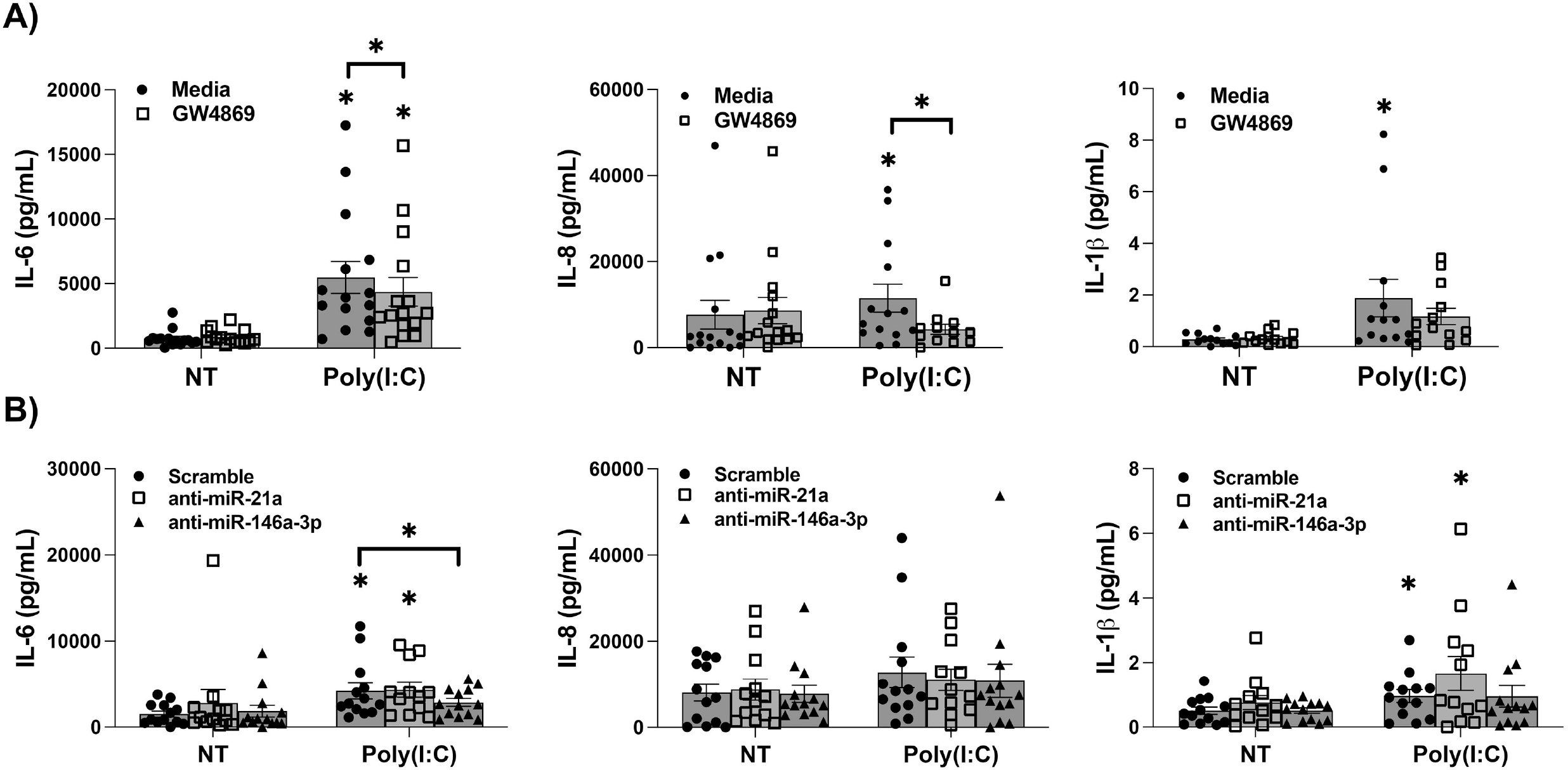
FM Poly(I:C)-induced pro-inflammatory factor release is partially dependent on sEVs and miR-146a-3p. (A) FM explants were pretreated with media or with the inhibitor of sEV biogenesis/release, GW4869, for one hour, and then treated with NT or Poly(I:C). After 24 hours, supernatants were measured for IL-6, IL-8 and IL-1β. (n=18-21) ^*^*p*<0.05 compared to the NT media control unless otherwise specified. (B) FM explants were transfected with either a scramble control, an anti-miR-146a-3p inhibitor, or an anti-miR-21a inhibitor for 24 hours and then treated with NT or Poly(I:C) for another 24 hours after which supernatants were measured for IL-6, IL-8 and IL-1β (n=13) ^*^*p*<0.05 compared to the NT control for each group unless otherwise specified.

### Poly(I:C)-induced FM secretion of IL-6 is dependent on miR-146a-3p

The role for TLR7/8-activating miRs was examined by transfecting FM tissues with specific miR inhibitors as previously reported^16^. The presence of the anti-miR-146a-3p inhibitor significantly reduced the ability of Poly(I:C) to trigger FM IL-6 secretion by 25.0±10.3%, while IL-8 and IL-1β secretion were unchanged (Figure 3B). The anti-miR-21a inhibitor did not significantly alter FM secretion of IL-6, IL-8, or IL-1β in response to Poly(I:C) when compared to scramble controls (Figure 3B).

## Discussion

Bacterial infections are well established as contributors to chorioamnionitis and preterm birth, a major threat to the health of the mother and child. Viral infections are also known to be contributors to chorioamnionitis and preterm birth; however, studies on the link and mechanisms between viral infections and preterm birth are lacking. While most viral research has focused on TORCH pathogens, less is known about non-vertically transmitted viruses that can still result in inflammation and preterm birth, such as influenza and SARS-CoV-2^4^. Previous studies using bacterial models of chorioamnionitis revealed that in response to PDG or LPS, miR-146a-3p is elevated in FM tissues and propagates inflammation in a TLR8-dependent manner both *in vitro* and *in vivo*^12,16^. In the current study, we investigated the role of this and other TLR7/8-activating miRs as intermediate regulators of FM inflammation in response to Poly(I:C), a viral dsRNA mimic.

In this current study we measured tissue expressed TLR7/8-activing miRs in FM explants. Surprisingly, and in contrast to our bacterial models^12,16^, none of the four miRs screened were increased, and miR-29a expression was significantly decreased following exposure to Poly(I:C). Nonetheless, we found that Poly(I:C)-induced inflammatory cytokine and chemokine secretion was in part dependent on TLR7 and TLR8 activation. In particular, as we have previously reported^12,16^, FM TLR8 had a role in both IL-6 and IL-8 production. Interestingly, in contrast to the bacterial model, TLR7 also played a role in mediating FM IL-8 secretion in response to Poly(I:C). While our data was not quite significant, both TLR7 and TLR8 may also play a role in regulating FM IL-1β secretion in response to Poly(I:C). Having established that both ssRNA sensors, TLR7 and TLR8, were playing a role in mediating downstream FM inflammation that was initiated by TLR3 activation in response to Poly(I:C), an alternative route of TLR7/8-activating miR delivery was investigated.

In pregnancy, sEVs serve as critical mediators of maternal physiological and immune changes as the pregnancy progresses. While sEVs have important functions to facilitate fetal development and the maintenance of pregnancy, they are also contributors to pregnancy complications such as preeclampsia and preterm birth^25^. Animal studies have shown that sEVs are a method of maternal-fetal communication and can induce preterm birth, hypothesized to be through inflammatory and oxidative stress signaling^26-28^. In a mix of cellular and animal studies, the importance of sEVs in maternal-fetal inflammation and the robust ability for sEVs to influence preterm birth is clear^26,29-31^. Despite this, identification of sEV cargo consistently involved in maternal-fetal inflammation and their specific pathways have remained elusive, especially in viral models. With the lack of an elevation in locally tissue produced TLR7/8-activating miRs in FMs treated with Poly(I:C), we investigated sEVs as a source of these DAMPs. Small EVs (sEVs) were isolated from FM conditioned media and confirmed via Western Blot for the positive markers, CD9 and CD81, as well as by nanoparticle tracking analysis. TLR7/8-activating miRs, miR-146a-3p and miR-21a, were found to be elevated in sEVs released from FMs exposed to Poly(I:C). Thus, we hypothesized that FM-derived sEVs provide an important mode of communication in viral TLR3-mediated chorioamnionitis and possibly in fetal inflammation since these sEVs can be released into the amniotic fluid. Indeed, TLR7/8-activating miRs are known to be packaged in sEVs and can exert stimulatory and inflammatory effects on target cells^18,19,22^. Furthermore, in a study measuring sEV miRs in preterm neonates, the group found that in neonatal bronchopulmonary dysplasia, sEV miR-21 was elevated with potential pro-inflammatory effects^32^. While FM inflammation was not studied alongside neonatal inflammation, the presence of miR-21 in fetal sEVs suggests a potential method of inflammatory communication between maternal and fetal tissues.

We predicted that in our current study, FM-derived sEVs were acting back in an autocrine manner to deliver TLR7/8-aciving miRs in order to propagate the inflammatory response to viral Poly(I:C). Thus, first to investigate the dependency of Poly(I:C)-induced FM inflammation on sEVs, we utilized the sEV biogenesis/release inhibitor, GW4869. GW4869 reduced the ability of Poly(I:C) to elevate FM secretion of both IL-6 and IL-8 in response to Poly(I:C). Interestingly, with sEV inhibition, Poly(I:C) induced IL-8 was reduced to baseline levels suggesting that IL-8 is mostly dependent on sEVs, while the IL-6 and potentially the IL-1β response may be only partially dependent upon this mechanism. Next, to investigate the dependency of Poly(I:C)-induced FM inflammation on the TLR7/8-activating miRs themselves, FM tissues were transfected with specific miR inhibitors. We found, somewhat surprisingly, that only miR-146a-3p appeared to regulate the FM response to Poly(I:C), and only the IL-6 response. It is important to note that in these experiments, FM IL-8 secretion was not significantly increased with Poly(I:C) treatment in the scramble controls when compared to NT scramble controls, suggesting that the transfections may have altered or dampened the IL-8 response. While miR-146a-3p and sEVs have direct effects on FM inflammation, in our model, miR-21a may be produced in response to an inflammatory stimuli, but may have a role independent of the generation of inflammation. Indeed, studies found that sEV miR-21a had cardio and lung-protective roles, and it was hypothesized to downregulate expression of pro-apoptotic/necroptotic genes^33,34^. Alternatively, it is possible that sEV miR-21a does have a pro-inflammatory role on factors or tissues that were not measured in the current project. Several studies suggest that sEV miR-21a have roles in intestinal sepsis^35^, macrophage polarization^36,37^, and neuroinflammation^38^, highlighting pro-inflammatory roles for sEV miR-21a. Further studies are needed to elucidate the role of FM-derived sEV miR-21a. It is also possible that the FM-derived sEVs under Poly(I:C) conditions contain other TLR7/8-activating DAMPs that we have yet to identify.

Throughout this project, IL-1β was limited in response to Poly(I:C) stimulation. While IL-1β may play a small role in TLR3-mediated inflammation, the anti-viral response may be more dependent on other factors such as IL-6 and IL-8, and possibly other factors not tested. Alternatively, in optimizing our studies for IL-6 and IL-8, we may not have captured the best timepoint for IL-1β secretion. Further limitations of this study include the variable nature human FM tissues and of Poly(I:C) as a TLR3 agonist. TLR3 recognizes dsRNA viruses; however, dsRNA viruses are limited in nature. More frequently, TLR3 senses dsRNA intermediates generated during the replication of ssRNA viruses^39^. This, in turn, may make TLR3 a robust secondary signaling mechanism, but a weak initial signal. Despite this, Poly(I:C) is a good viral mimic to test our hypothesis of miRs acting as DAMPs and secondary signals via TLR7 and TLR8 in FM inflammation.

With ongoing research efforts, the effects of viral infections on pregnancy outcomes are becoming clearer. Despite this, the mechanisms involved in the maternal viral response and their role in preterm birth is not well understood. Herein, we have demonstrated that TLR7/8-activating miR-146a-3p is an important pro-inflammatory DAMP mediating FM responses to a viral stimuli. We propose that upon recognition of a virus, FM miR-146a-3p is elevated and packaged into sEVs for delivery to neighboring cells where it perpetuates inflammation via TLR7 and TLR8 activation in and autocrine and paracrine manner within the tissue. Identification of this novel anti-viral signaling mechanism provides an opportunity for further exploration and a possible diagnostic or intervention step to mitigate inflammation induced preterm birth.

## Acknowledgements

The authors would like to thank the Yale University Reproductive Sciences Biobank and the staff of Labor and Delivery for tissue collection. This research made use of the Yale University Biophysical Resource Core for Nanoparticle Tracking Analysis.

